# Comparison of oviposition and adult trapping to monitor *w*Mel introgression for *Wolbachia-*based vector control

**DOI:** 10.1101/2025.03.10.642347

**Authors:** Elisabeth Nelson, Thiago N. Pereira, Erica Milena de Castro Ribeiro, Bianca Daoud Mafra e Silva, Carolina Camillo, Thiago Rodrigues da Costa, Mauro M. Teixeira, Albert I. Ko, Derek A. T. Cummings, Luciano A. Moreira

## Abstract

*Wolbachia* introgression into *Aedes aegypti* mosquito populations has been shown to be effective in preventing dengue and is being evaluated for WHO prequalification. Monitoring the long-term introgression of *Wolbachia* (*w*Mel)-positive *Aedes aegypti* mosquitoes, however, requires labor-intensive and costly BG-Sentinel traps (BG-traps). More affordable alternatives, such as using oviposition traps (ovitraps), have not been fully evaluated.

*Ae. aegypti* eggs and adults were collected from 124 ovitraps and 237 BG-traps, respectively, across 12 clusters in Belo Horizonte, Brazil from March to May 2023 as part of the EVITA Dengue trial. We used a qPCR assay to detect *w*Mel in a sample of L3-L4 stage larvae (up to 29) that were reared from eggs in ovitraps and adults from BG-traps (up to 10 per BG-trap). We used mixed effects models to compare estimates of cluster-level *w*Mel introgression from ovitrap and BG-trap data over time.

Among 3,675 larvae reared from ovitraps, *w*Mel prevalence was 0.50 (95% CI: 0.48-0.51). Among 1,244 adult *Ae. aegypti* tested from BG-traps, *w*Mel prevalence was 0.45 (95% CI: 0.42-0.48). Cluster-level *w*Mel introgression in larvae and adults was highly correlated (Spearman’s r = 0.70, p = 6.71e-06). Multivariate analysis found that ovitrap estimates of introgression were associated with BG-trap estimates in the same month when models incorporated the previous month’s ovitrap *w*Mel-positive count, the proportion of *w*Mel in ovitraps in the current and previous month, and *Ae. aegypti* abundance. Leveraging this model, predicted *w*Mel introgression from ovitrap data were highly correlated with observed introgression from BG-trap data (r_s,counts_=0.98, p=1.53e-14; r_s,prevalences_=0.82, p=0.11e-05) and provided greater precision than crude ovitrap-based estimates.

These findings indicate that ovitrap-based monitoring represents a low cost, more efficient approach to evaluating introgression as the *Wolbachia*-based interventions are scaled up and implemented broadly in high burden regions for dengue and other arboviral diseases.

**Author Summary:** Dengue fever is a major global health burden, and one promising way to control it is by releasing *Aedes aegypti* mosquitoes infected with a bacteria called *Wolbachia*. This bacteria reduces the mosquitoes’ ability to spread dengue. However, monitoring the success of *Wolbachia* in mosquito populations over time requires expensive and labor-intensive traps.

Our study explored a more affordable alternative: using oviposition traps (ovitraps), which collect mosquito eggs instead of adults. In Belo Horizonte, Brazil, we compared data from eggs collected in ovitraps with data from standard mosquito traps (BG-traps) that catch adult mosquitoes. By analyzing the mosquitoes for *Wolbachia*, we found that the egg-based method provided reliable estimates of *Wolbachia* levels in the mosquito population.

These results suggest that ovitraps could be a cost-effective and efficient way to monitor *Wolbachia*’s spread. This approach could help improve dengue prevention efforts, making it easier for public health programs to track and expand this control strategy in areas where dengue is a major concern.

## Introduction

*Wolbachia* introgression into *Aedes aegypti* mosquito populations has been shown to be highly effective in preventing dengue. In the first cluster randomized-controlled trial of this intervention in Yogyakarta, Indonesia, releases of *Wolbachia* (*w*Mel)*-*infected *Ae. aegypti* reduced virologically-confirmed dengue by 77.1%, despite widespread contamination between intervention and control clusters (1). In large-scale releases in Rio de Janeiro, Brazil, increasing reduction in dengue incidence was associated with increasing *Wolbachia* introgression (2). Reductions in dengue incidence of 71% were observed in areas where >60% *Wolbachia* introgression had occurred. Yet, decreased dengue risk of 20% was also seen in areas where *Wolbachia* introgression was low (10-20%). Although evidence is accruing that demonstrates the effectiveness of *Wolbachia*-based interventions, scaling these interventions for widespread roll-out in low resource settings will be a key challenge in the future.

Monitoring *Wolbachia* introgression during and after release of *w*Mel-infected *Ae. aegypti* is a key implementation barrier. Surveillance on introgression and wildtype *Ae. aegypti* populations is used to guide releases to achieve introgression. Modeling studies have suggested that once introgression levels are greater than 60%, local spread of *Wolbachia* is highly likely to increase due to the frequency-dependent fitness advantage of *Wolbachia* infected females (3). Studies also suggest that *Wolbachia* introgression levels greater than 60% lead to a self-sustaining *Wolbachia* populations (3). Therefore, it is crucial to accurately estimate *Wolbachia* introgression to ensure successful, self-sustaining introgression into wildtype populations and successful prevention of arboviral transmission. Current methods require an extensive network of mosquito traps to continuously monitor local mosquito populations. BG-Sentinel traps (BG-traps) are the standard for monitoring *Wolbachia-*based interventions, but they are expensive and require a constant source of power which in turn restricts where they can be deployed (4,5).

Oviposition traps (ovitraps) present a potential alternative monitoring method to help overcome the cost and resource barriers to scale-up. However, there is limited information on the accuracy of ovitraps’ prediction of *Wolbachia* introgression. *de Jesus et al*. conducted a proof-of-concept study using primarily semi-field experiments and a small field comparison during ongoing releases in Rio de Janeiro to compare the efficacy of ovitraps to BG-traps (4). They found a low (<0.1%) sampling error using ovitraps in a semi-field enclosure and comparable introgression predictive abilities between ovitraps and BG-traps in a small field-based experiment. Despite these results, the predictive ability of ovitraps at estimating *Wolbachia* introgression has yet to be fully evaluated in field settings and at larger scales.

We had the opportunity to address this question during the implementation of the EVITA Dengue trial, a cluster-randomized controlled trial that is evaluating the effectiveness of releasing wMel-infected *Aedes aegypti* in preventing arboviral infection in the city of Belo Horizonte, Brazil (6). As part of this trial, BG-trapping was performed in intervention and control clusters, in parallel to monitoring ovitraps which were performed by local public health authorities. Our study aimed to correlate *w*Mel prevalence estimates measured using ovitraps to those measured using BG-traps and evaluate the accuracy of ovitrap-based estimates of introgression in predicting BG-trap estimates.

## Methods

### EVITA Dengue Trial

Our study was conducted in Belo Horizonte, the sixth largest city in Brazil, located in the Southeastern state of Minas Gerais. Located at 850m above sea level, the climate is primarily tropical, receiving around 50 inches of rainfall a year, with an arid winter season (7). This city was selected as the site for EVITA Dengue trial, a cluster-randomized control trial that is evaluating the effectiveness of releasing *w*Mel *Ae. aegypti* adults on the risk of serologically-ascertained arboviral infection among schoolchildren, which is scheduled to end in December 2024 (6). Within the city, 58 school-based clusters were identified, of which 29 were randomized to intervention and 29 control arms. Clusters were 0.44 – 2km^2^ in size and separated by at least 200m or a natural barrier boundary from another cluster in order to mitigate the risk of contamination of control clusters with *w*Mel *Ae. aegypti* from intervention clusters.

The nine major municipal regions of the city were geographically-grouped into three phases of staggered releases, to improve operational feasibility: Phase 1 underwent releases between January 2021 – January 2022 in 10 clusters, Phase 2 underwent releases during April 2021 – April 2022 in 9 clusters, and Phase 3 had releases between May 2021-May 2022 in 10 clusters. Intervention and control clusters were balanced within each of these Phases. Detailed information on the releases and intervention monitoring can be found in the *Collins et al.,* the trial protocol (6).

### Study Sites

Nine intervention clusters, one from each of the city’s main areas (three from each of the trial release phases), and three control clusters, one from each of the trial release phases, were selected by an unblinded member of the World Mosquito Program (WMP) team and assigned a letter to ensure continued blinding. Samples were collected from March – May 2023 (epidemiological weeks 10, 14, and 18) during the monitoring phase of the EVITA trial. Sample processing then occurred between mid-May – late July 2023. Since *w*Mel-positive eggs do not remain viable when desiccated as long as wildtype (*w*Mel-negative eggs) it was determined that March was the earliest collections could begin to be stored prior to May processing (8).

### BG-Sentinel Trapping

The BG-Sentinel trap, which is currently the gold-standard for capturing *Ae. aegypti* mosquitoes, uses visual, olfactory, and chemical attractants to mimic convection currents created by the human body to capture host-seeking females (4,9,10). As part of the EVITA trial, these traps were placed in a gridded arrangement in all clusters, with up to 16 traps per km^2^ (6). Traps were preferentially placed in commercial locations, such as stores, one meter or lower from the ground. The traps used in the trial do not use any chemical bait or CO_2_, instead only attracting females through their dark color (11). Given that our study took place during the monitoring phase of the EVITA trial, traps operated for 15 days per month and samples were collected monthly. Trap collections were processed at the WMP Biofactory in Belo Horizonte, where all mosquitoes were morphologically identified at the species level and recorded. Up to 10 *Ae. aegypti* samples per trap were then sent to the WMP Diagnostic Laboratory at the Instituto René Rachou (Fiocruz) for *w*Mel identification (12).

### Oviposition Trapping

Oviposition traps consist of a wooden paddle submerged in a container of hay-infused water (13). Belo Horizonte’s municipal government has maintained a surveillance network of ovitraps, placed at intervals of 400m, monitored for arboviruses and mosquito abundances monthly since 2006. The traps were installed outside of buildings. Staff of the Municipal Ministry of Health checks traps bi-weekly, with traps installed for seven days every two weeks in an alternating pattern around the city. There were 5-15 ovitraps per cluster, which have been found to collect on average, 22-42 eggs per trapping cycle.

The Ministry of Health delivered ovitrap paddles from the study clusters to the WMP Biofactory, where they were sorted by cluster and time point. Eggs were hatched in a controlled environment at 28±2°C, 82%±2 relative humidity and 12h light/dark regime. Larvae were fed a liquid solution of liver and yeast paste. L3-L4 stage larvae, the third and fourth instar phases of larval development, samples were collected and taken to WMP Diagnostic Laboratory for *w*Mel identification.

The minimum sample size was determined to be approximately 22 *Ae. aegypti* mosquitoes per ovitrap per time point to measure *Wolbachia* prevalence with 80% power. This was calculated using a two-sided power calculation for proportion test (one sample) from the ‘pwr’ package in R, assuming *w*Mel prevalence was at least 60% in all intervention clusters, 80% power, and 5% error (Supplemental Fig S1). Given the ovitrap collections generally contain approximately 10% *Aedes albopictus*, 28-29 larvae were taken from each ovitrap sample, to try to ensure 22 *Ae. aegypti* samples. If 28-29 larvae were not available for a collection, the total number of larvae available were used.

### Estimation of wMel Prevalence

All samples were processed using WMP’s standardized protocols, which have been validated throughout their international study sites (12). For DNA extraction, adults and larvae were individually macerated and homogenized in Squash Buffer Mix (pH 8.2, 10 mM Tris–Cl, 25 mM NaCl, 1 mM EDTA), supplemented with Proteinase K (200 µg/ml), followed by incubation at 56°C for 5 min to activate the enzyme. Extraction was interrupted by enzyme inactivation at 98°C for 5 min. The plate was then cooled to 12°C.

For *w*Mel identification, adults and larvae were screened using the Promega qPCR kit, on a QuantStudio12k. The qPCR cycling program consisted of a preincubation at 95°C for 5 min, followed by 40 cycles of PCR amplification (95°C for 3 seconds and 60°C for 30 seconds), and a cooling down step at 37°C for 30 seconds. *Ae. aegypti* and *w*Mel were identified using the endogenous control housekeeping gene RPS17 for *Ae. aegypti* and the gene encoding *Wolbachia* surface protein WSPTM2 using the following primers and probes: RPS17 Forward: 5′-TCC GTG GTA TCT CCA TCA AGC T-3′, Reverse: 5′-CAC TTC CGG CAC GTA GTT GTC-3′ and Probe 5′-HEX/CAGGAGGAG/ZEN/GAACGTGAGCGCAG/ 3lABkFQ-3′; WSPTM2 Forward: 5′-CATTGGTGTTGGTGTTGGTG-3′ and Reverse: 5′-ACACCAGCTTTTACTTGACCAG-3′ and WSPTM2 Probe 5′-FAM/TCCTTTGGA/ ZEN/ACCCGCTGTGAATGA/3lAbRQSp-3′. Individual mosquitoes were determined to be positive for *w*Mel if the qPCR Ct value of WSP was below 29 and determined to be an *Ae. aegypti* specimen if the qPCR Ct value of RPS was below 28, whereas *Aedes albopictus* were identified if the qPCR Ct value of the RPS was above 28.

Individual trap-level prevalence of *w*Mel-positive *Ae. aegypti* out of total *Ae. aegypti* tested were averaged by cluster and time-point to calculate cluster-level introgression estimates. We excluded data from ovitrap paddles that did not result in any hatched larvae (n = 178), as this may have been due to paddles not being collected by the municipality at that time point or eggs falling off in transport.

### Statistical Analyses

Statistical analyses were performed in R, version 4.2.2 (14). An initial analysis to determine whether a significant difference existed between the introgression estimates of each trap type was performed using a paired t-test. Prevalences by cluster and time-point were paired between BG-traps and ovitraps. The correlation between the trap-type estimates was then tested using a Spearman-Rank Correlation Test, which measures the strength and direction of association between ranked variables and is robust to non-linearity. We plotted the cluster-level prevalence estimates by time point of BG-traps (x-axis) and ovitraps (y-axis). We then determined the sensitivity and specificity of the ovitrap-estimated prevalences by cluster and time point in classifying prevalence estimates as meeting an introgression threshold of 60%, with BG-trap based prevalence providing the gold standard. We also investigated a threshold of 50%. Additionally, we examined the within-cluster variance in estimated *w*Mel prevalences by trap type and time point visually, using the *ggplot2* package in R to construct forest plots.

To evaluate predictions of BG-trap results using ovitrap measurements, we constructed negative binomial generalized linear mixed-effects regression models, with the outcome variable of BG-trap estimated *w*Mel introgression. Potential predictor and covariate variables included: ovitrap estimated *w*Mel count, ovitrap estimated *w*Mel proportion, lags of ovitrap count and proportion, month, an interaction between ovitrap proportion and month, and cluster. Given that ovitrapping targets a part of the mosquito life cycle that is shifted in time from the adult phase (either forward or backward in time), we explored the correlation of multiple lags of counts from each trap type using a Spearman-Rank Correlation test. Given the constrained time scale of the data, three monthly observations per trap type per cluster, only lags of ±1 and zero months were used. The variable with the highest correlation was then used as the primary predictor in the models.

All models were fit with an offset of the log of total *Aedes* caught in BG-traps in that cluster at that time point. An offset was used, as this renders the outcome effectively as a rate and allowed us to include uncertainty around estimates in traps with fewer total mosquitoes caught. All combinations of covariates were tested using cluster-level data, to maintain blinding for the *EVITA Dengue* trial. Models were constructed using the *glmer.nb* function in the *MASS* package in R. Model building and selection took place in two phases: 1) examining the relationship between BG-trap and ovitrap data by month, cluster, and covariate space; 2) using the best fitting model for prediction to determine whether ovitrap data is a good proxy for BG-trap data. Model selection in the second phase was determined using the Bayesian Information Criterion score (BIC) using all possible combinations of covariates. The predicted *w*Mel counts and introgression estimates from the best fitting model were compared to the real BG-trap counts and introgression estimates using a Spearman-rank correlation test and sensitivity and specificity tests of the predicted *w*Mel counts or introgression estimates. Finally, the magnitude and directionality of the model’s error was investigated by comparing the predicted *w*Mel counts to the real BG-trap counts. Error was calculated by subtracting the BG-trap count from the model predicted count and dividing by the BG-trap count.

## Results

### wMel Prevalence in BG-Traps and Ovitraps

During epidemiological weeks 10, 14, and 18 in March, April, and May of 2023, 1,580 *Ae. aegypti* and *Ae. albopictus* adults were captured by BG-traps in the 12 selected clusters in Belo Horizonte. Of these, we tested 1,244 adult *Ae. aegypti*, with an overall *w*Mel prevalence of 0.45 (0.42-0.48) (Fig 2A-C shows cluster level proportions). At these same time points, 6,919 larvae hatched and were reared from ovitraps, with an overall hatch rate of 45% (6,919/15,242). The hatch rates were lower than expected, which have been found to normally be around 80%, with the highest rate observed in March (52%) and then similar rate in April (42%) and May (46%) (15–17). These lower-than-expected hatch rates were likely due to the increasingly cold temperatures with the onset of winter or handling and storage of the ovitrap paddles. We tested 3,675 larvae, of which 3,149 were *Ae. aegypti* (86%) and 609 were *Ae. albopictus* (17%), with an overall *w*Mel prevalence of 0.50 (0.48-0.51) among *Ae. aegypti* tested (Fig 2A-C). *w*Mel prevalence estimates appeared to follow a similar pattern across clusters and time points between the two trap types (Fig 2A-C).

**Fig 1.**
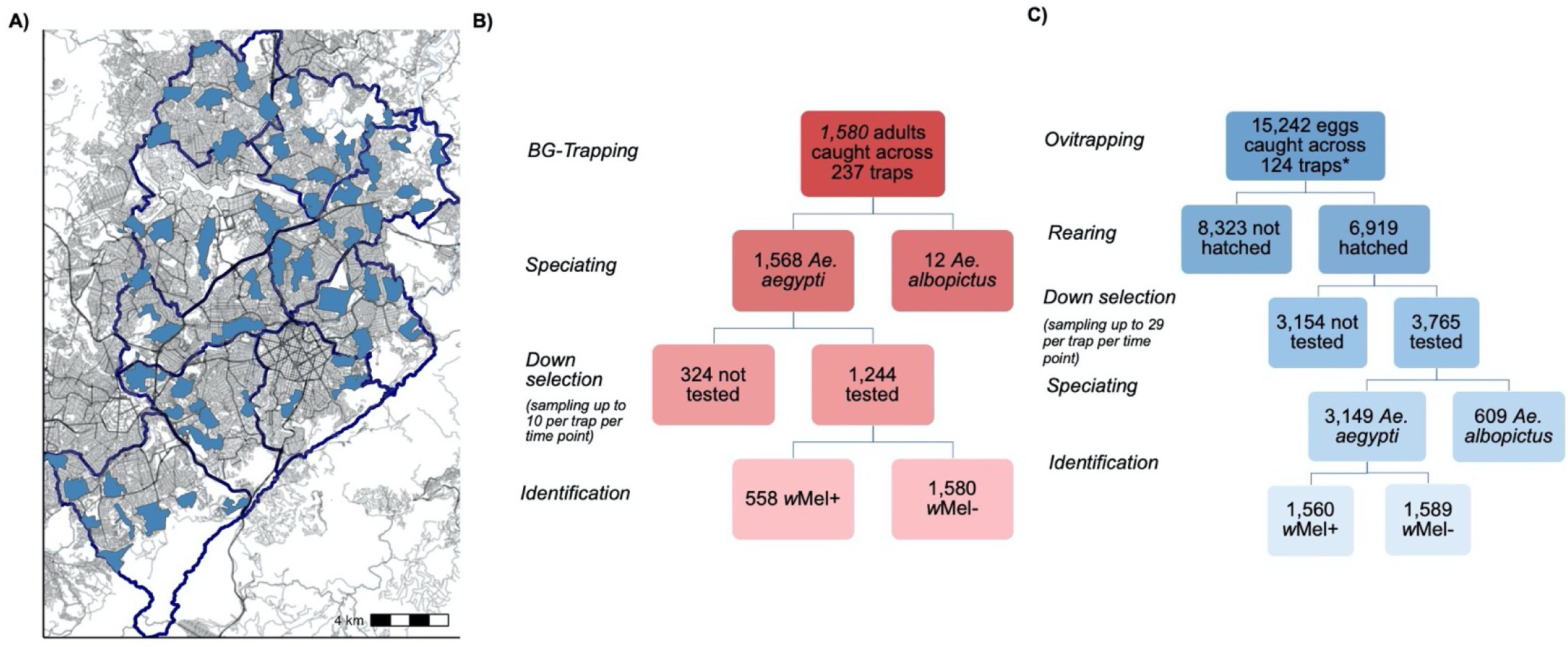
Experimental Study design. A) EVITA Trial clusters in Belo Horizonte, Brazil with road density and size indicated by grayscale (6). B) *Aedes a*dult mosquito testing protocol from BG-trap collection. C) Larvae mosquito testing protocol from ovitrap collection. Down selection refers to the sampling of trap collections for testing: up to 10 *Ae. aegypti* adults per BG-trap and up to 29 *Aedes* larvae per ovitrap. *Eggs were counted by the Municipal Ministry of Health in March and April and by the first author in May.

**Fig 2.**
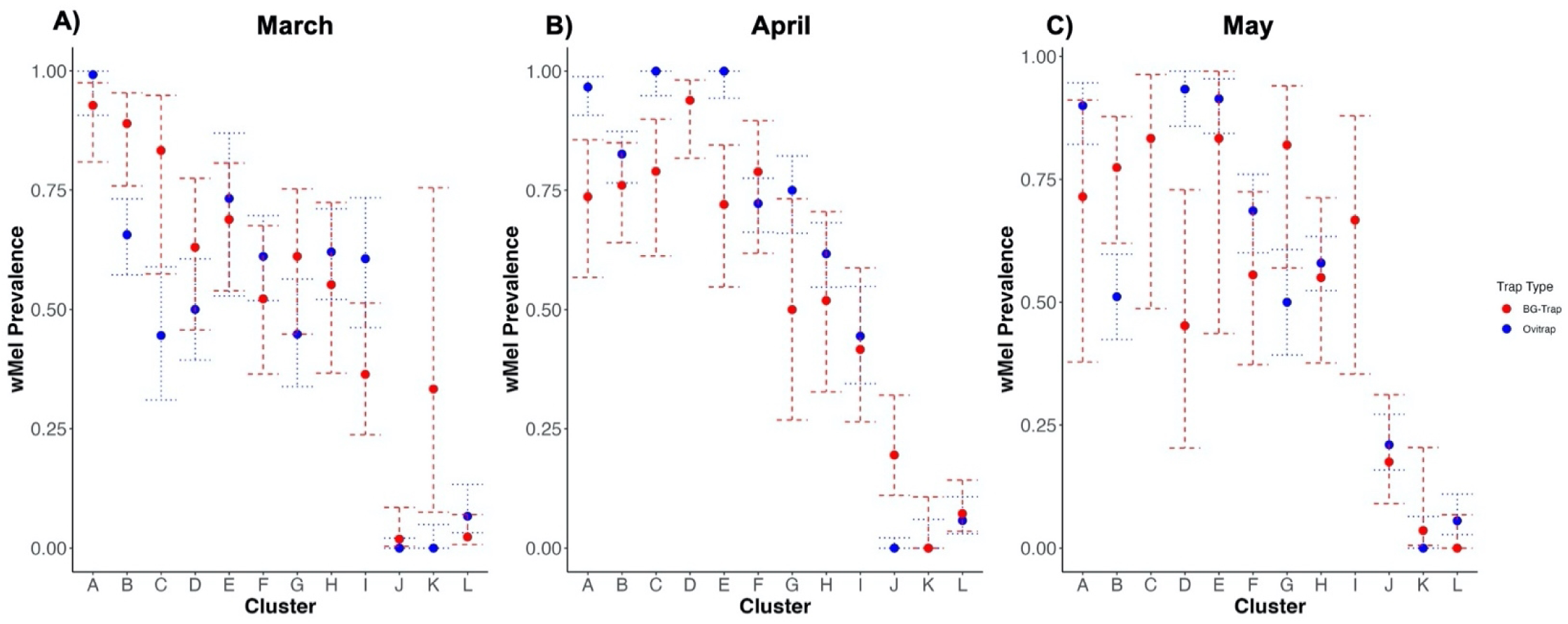
*w*Mel prevalence estimates by cluster and trap-type March-May 2023. *w*Mel prevalence estimates in adult mosquitoes from BG-traps and ovitraps in March (A), April (B), and May (C). The point estimate is surrounded by its 95% binomial confidence interval. The cluster-level prevalence estimates represent an average of the trap-level proportions in that cluster at that time point.

**Fig 3.**
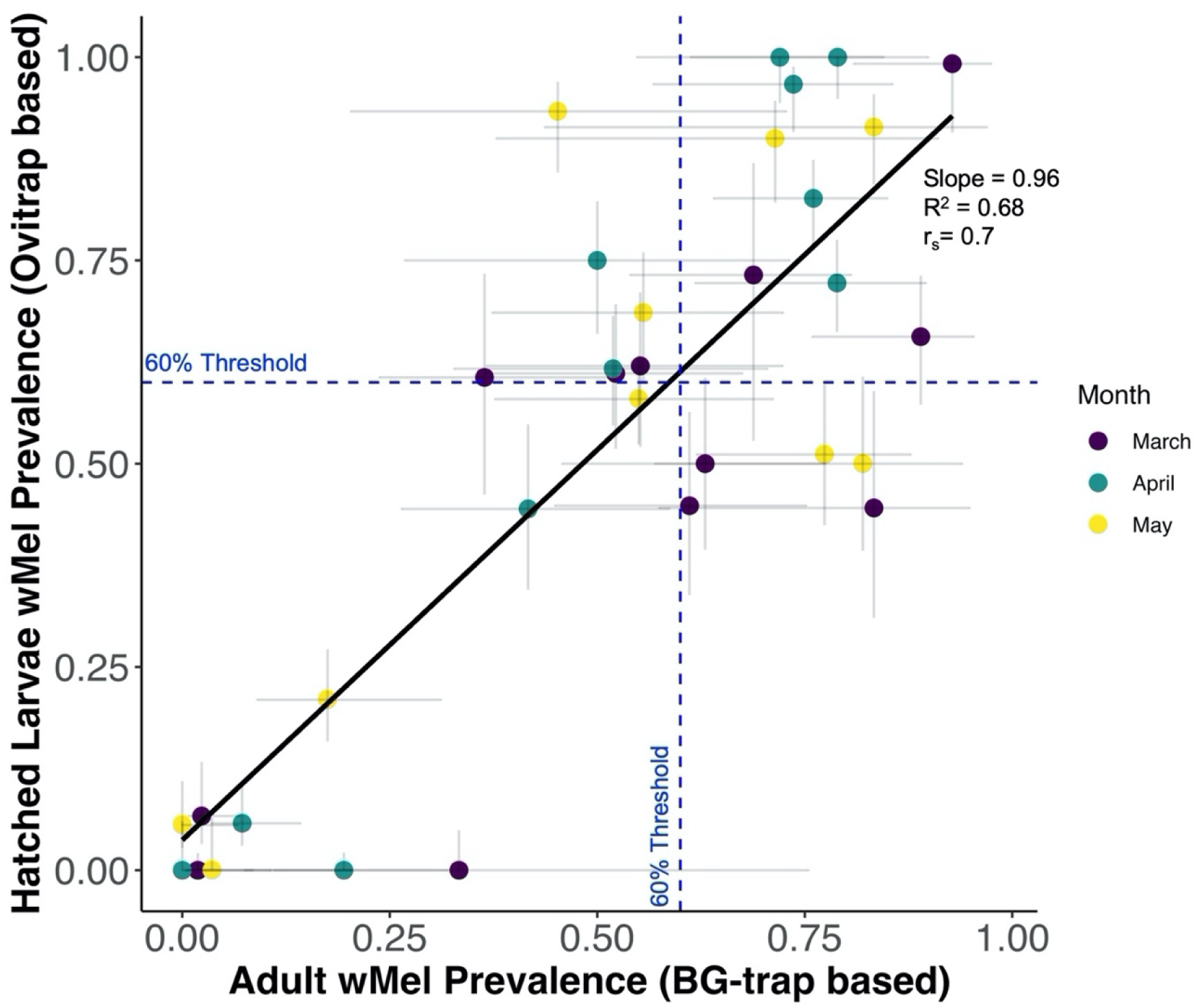
Correlation of cluster-level *w*Mel prevalence estimates in adults and larvae. The correlation between *w*Mel prevalence estimates from *Ae. aegypti* adults collected in BG-trap and larvae hatched from ovitrap collections by cluster and time point. The color of the point estimate indicates the time point, with purple for March, blue for April, and yellow for May. In the x-direction 95% binomial confidence intervals from the BG-trap estimate extend from the point, while in the y-direction 95% binomial confidence intervals from the ovitrap estimate extend from the point. The black line depicts the linear relationship between the BG-trap adult prevalence estimates and the ovitrap hatched larvae prevalence estimates, calculated using a linear model, with corresponding linear slope, adjusted R^2^, and Spearman-rank correlation. The blue dotted lines indicate a 0.6 (60%) *w*Mel introgression threshold.

### Correlation between wMel Prevalences in BG-Traps and Ovitraps

We did not find significant difference between the ovitrap and BG-trap cluster-level *w*Mel prevalence estimates at each time point (mean difference = 0.002, 95% CI = −0.05 – 0.09, p=0.62), using a paired t-test. Spearman-rank correlation tests found high correlation between ovitrap and BG-trap cluster-level estimates of *w*Mel prevalence at each time point (r_s_=0.70, p=6.71e-06, R^2=0.68), suggesting a strong association of ranks. Ovitraps correctly classified the introgression threshold in 64% of estimates. However, 12 estimates (36%) were misclassified, with 7 overestimated by ovitraps and 5 underestimated. The overall sensitivity of ovitrap prevalence classification was 59% and the specificity was 69%, across clusters and time points. The sensitivity and specificity appeared to be the lowest in March, the month of the greatest number of misclassifications as well (Supplemental Table S5). When the introgression threshold was lowered to 50%, only four misclassifications occurred. If a further reduced threshold of 36% was used, there was no discordance between the two trap types’ estimates.

Examining the within-cluster variation of prevalence estimates, i.e. the variance in the trap-level estimates, we found similar within cluster variation between ovitraps (0.13) and BG-traps (0.12), across clusters and time points. When viewed by time point, ovitraps had slightly lower within-cluster variance in April and May (0.17 in March, 0.09 in April, 0.11 in May) than BG-traps (0.11 in March, 0.13 in April, 0.13 in May), however not in March (Supplemental Fig S3).

### Factors Associated with BG-trap-based wMel Introgression

To determine the best fitting model of the *w*Mel-positive *Ae. aegypti* captured by BG-Traps (outcome) and *w*Mel-positive *Ae. aegypti* captured by ovitraps (predictor), we tested the correlation by cluster of ±1 month temporal lags of ovitrap and BG-trap *w*Mel-positive mosquito counts, using a spearman rank correlation test (Supplemental Table S6). We found a slightly positive correlation between BG-trap and ovitrap counts averaged across locations through time (r_s_=0.12). We found a strongly positive correlation between BG-trap counts and a −1 month lagged ovitrap count (r_s_=0.78), as well as between a +1 month lagged BG-trap count and ovitrap count (r_s_=0.78). The *Ae. aegypti* life cycle takes around 7-10 days to complete and the adult life span is approximately three weeks, so eggs one month would likely be adults the next and vice-versa. Therefore, it is both statistically- and biologically-plausible that there is a lagged temporal correlation between BG-trap and ovitrap *w*Mel-positive counts. Given that our model aimed at investigating the association with the current month’s BG-trap *w*Mel count, we used a −1 month lagged ovitrap *w*Mel count as the primary predictor variable in all models.

Using a negative binomial regression approach we examined the relationship between BG-trap and ovitrap data by month, cluster, and covariate space. Across models, the average coefficient for the −1 month lagged ovitrap *w*Mel count was −0.00057. Given that the outcome of the model was a proportion (introgression), an incremental increase in the number of *w*Mel positive larvae in an ovitrap would not likely affect the overall prevalence of *w*Mel in the cluster, and therefore a coefficient of almost zero is reasonable. The prevalence of *w*Mel in ovitraps had an average coefficient of 1.178 across models, showing a significant association with increases in the outcome proportion in BG-traps. The −1 month lagged ovitrap proportion showed greater variability across models, but with an average coefficient of 1.354. Therefore, it appears that the prevalence of *w*Mel in ovitraps in both the current month and the previous month were highly associated with the current month’s prevalence of *w*Mel in BG-traps, when accounting for the previous month’s ovitrap *w*Mel count. Finally, including an interaction between ovitrap prevalence and month did not appear to help the fit of the model greatly. While surprising, this reinforces the stability of the association between ovitrap prevalence and BG-trap prevalence through time.

### Predictive Model of BG-trap-based wMel Introgression using Ovitrap-based inputs

We found the best fitting model, determined by BIC scores, included the covariates of −1 month lagged ovitrap *w*Mel count, ovitrap *w*Mel prevalence, −1 month lagged ovitrap *w*Mel prevalence, and an offset of the total *Ae. aegypti* caught by BG-Traps (Model 5 in Table 1, bolded). Additionally, the model including only −1 month lagged ovitrap *w*Mel count, ovitrap *w*Mel prevalence, and an offset of the total *Ae. aegypti* caught by BG-Traps (Model 2 in Table 1) fit almost as well and may provide a simpler model for prediction by requiring slightly less information. Using these models, we compared the model prediction accuracy to the gold-standard approach of BG-traps in estimating *w*Mel counts and inrogression. As shown in Fig 4, the predicted *w*Mel counts were highly positively correlated with the true counts from BG-traps for both models (Model 5: r_s_=0.98, p=1.53e-14; Model 2: r_s_=0.93, p=4.15e-09). The model predicted introgression estimates were also highly positively correlated with the BG-trap introgression estimates (Model 5: r_s_=0.82, p=0.11e-05; Model 2: r_s_=0.64, p=0.0024), however model 2 was slightly less correlated than the raw ovitrap introgression estimates (r_s_=0.70). Overall, the model predicted the real *w*Mel counts very well, with an error of 0.0004 for Model 5 and 0.004 for Model 2 across clusters and time points (Supplemental Table S7). Both models slightly underestimated counts in April (Model 5 error = - 0.02; Model 2 error = −0.002) and slightly overestimated in May (Model 5 error = 0.02; Model 2 error = 0.01).

**Table 1.**
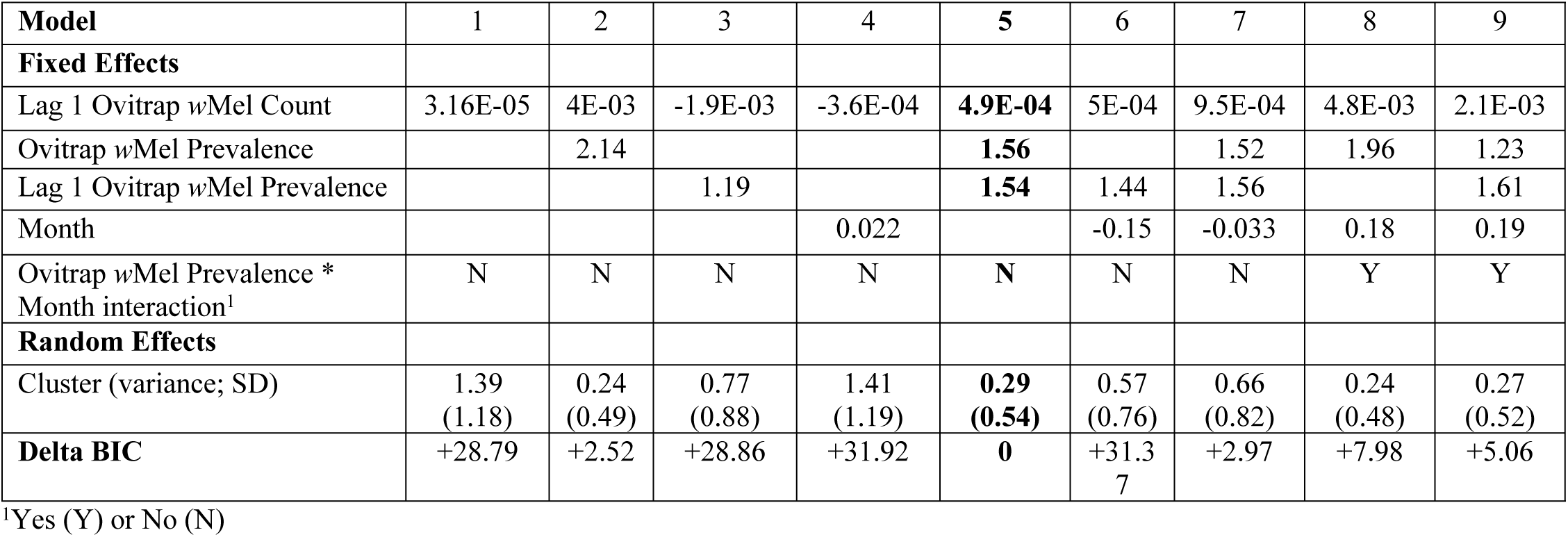
Regression model coefficients of a model predicting *Wolbachia* prevalence as measured with BG-traps. The relationship between ovitrap and BG-trap collected data was modeled using a negative binomial regression approach. Cluster was included as a random effect in all models as well as an offset using the log of total *Ae. aegypti* caught by that BG-trap. Bayesian Information Criteria (BIC) is presented as 0 for the best fitting model and the difference in BIC for the other models.

**Fig 4.**
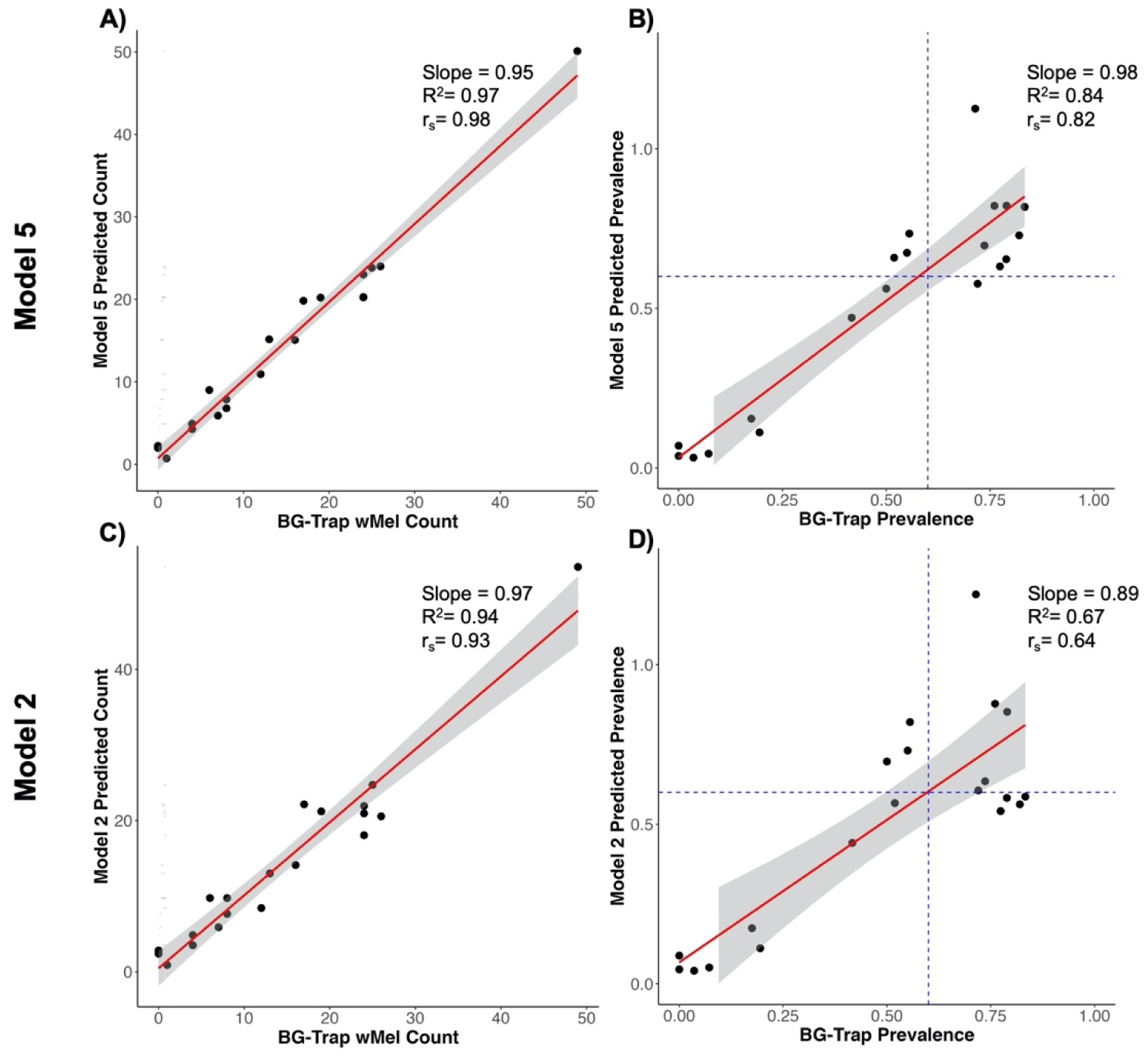
Model prediction. Correlation of BG-Trap and Model 5 predicted *w*Mel counts (A) and introgression estimates (B) by cluster and time point. Correlation of BG-Trap and Model 2 predicted *w*Mel counts (C) and introgression estimates (D) by cluster and time point. The red lines depict the linear relationship between the BG-Trap and the model predicted collections/estimates, calculated using a linear model, with corresponding slope, adjusted R^2^, and Spearman-rank correlation.

Utilizing the 60% WMP introgression threshold, we found the overall sensitivity of the Model 5’s introgression classification was 73% and the specificity was 89%, across clusters and time points, and Model 2’s sensitivity was 63% and specificity was 67% (Supplemental Table S8). Model 5 correctly classified 80% of the estimates, but misclassified 4 estimates (20%), with 3 overestimated and 1 underestimated. Model 2 correctly classified 65% of the estimates, but misclassified 7 estimates (35%), with 3 overestimated and 4 underestimated.

## Discussion

*w*Mel *Wolbachia* is poised to play a large role in the future of arboviral disease control given the mounting evidence of its effect on reducing arboviral infection disease risk. Resource constraints may hamper efforts to scale-up this intervention. In this study, we aimed to determine the predictive precision of a low-resource monitoring strategy, oviposition trap surveillance, in a post-release phase of a *w*Mel program compared to the more resource intensive BG-traps. Using the sampling strategy and cluster demarcations of the EVITA randomized control trial and municipal arboviral surveillance in Belo Horizonte, Brazil, we compared the introgression estimation of ovitraps and BG-traps. We have shown that data from ovitraps is highly correlated with that from BG-traps and serves as good proxy of *w*Mel-positive introgression estimates from BG-traps. However, given the limited number of months of evaluation, further work is needed to assess the generalizability of ovitrap results and their accuracy compared to BG-traps.

We found cluster-level *w*Mel introgression estimates were highly correlated between ovitraps and BG-traps over time. While the overall sensitivity and specificity of ovitrap introgression classification was moderate, 67% and 61% respectively, half of the misclassifications occurred in March. Eggs from March ovitrap paddles were stored until testing in early June, so the misclassification may be due in part to the proven lowered quiescent viability of *w*Mel-infected eggs. In other studies we performed on *w*Mel detection in eggs, we found that four weeks appears to be the longest eggs can be stored before the failure rate of detection significantly increases. Therefore, to avoid this misclassification in the future, eggs should not be stored for greater than four weeks before *w*Mel detection.

In our regression modeling analysis, we observed that by accounting for the previous month’s ovitrap *w*Mel-positive count, the prevalence of *w*Mel in ovitraps in both the current month and the previous month, and information on *Ae. aegypti* abundance, ovitrap estimates were highly associated with the current month’s count of *w*Mel-positive in BG-traps. In our best fitting predictive model (Model 5), we were able to leverage this association to estimate *w*Mel counts and prevalence estimates with greater precision than just using ovitrap introgression estimates alone. Given that our models do not include a monthly temporal term, their results are likely generalizable to other time periods if a measure of *Ae. aegypti* abundance is known (i.e. the offset we included in our models). With low error and relatively high sensitivity and specificity, our best fitting predictive model provides a framework for future use of ovitrap monitoring to determine *w*Mel introgression.

This study investigates the comparative predictive precision of ovitraps and BG-traps for the monitoring of *w*Mel introgression in a post-release monitoring phase of *w*Mel releases. A previous study investigated the potential use of ovitraps in monitoring *w*Mel during releases in Rio de Janeiro, however the study focused on a proof-of-concept approach by conducting a majority of lab-based and semi-field experiments, with a small-scale field data collection (4). They found a sampling error of less than 0.1% using ovitraps in their semi-field system and found similar predictive abilities between the ovitraps and BG-traps in the Urca neighborhood. Our study aimed to build upon the results of this previous study, expanding to a new geographic region, new season, with the added strength of the EVITA Trial cluster design, and utilizing a predictive modeling framework.

While there are numerous strengths to this study, there are certain inherent limitations as well. Given the ongoing nature of the EVITA Trial, no spatial information was available for the traps. While this maintained the blinding of the investigators, it limits the resolution at which associations can be made to the cluster-level. Further exploration is needed to determine the spatial correlation of BG-traps and ovitraps in close proximity and the resolution of prediction the trap network creates. However, by investigating the within-cluster variance of both trap types, we found consistently low variance for both, which suggests relatively low trap-to-trap heterogeneity within clusters. In tandem with the high autocorrelation of BG-trap estimates found in our sensitivity analysis, it is unlikely that trap positioning drives large trap-to-trap differences in prevalence estimates. Both ovitrap-derived and model-predicted prevalence estimates exhibited misclassification using a 60% introgression threshold, which may lead to operational challenges, as WMP gauges when to stop releases in the consolidation phase based on this threshold. However, the majority of estimates were correctly classified, and we hypothesize the misclassification may have mainly been due to the storage time of the eggs. In the future, storage of ovitrap paddles before rearing and testing should be limited. Additionally, in the release and consolidation phases of release programs, *w*Mel introgression is likely to be <40%, a threshold for which our models showed little misclassification. Finally, we were not able to perform out-of-sample validation of our models due to the limited temporal extent of our data. Future ovitrap data collection will allow us to externally validate the model.

## Conclusion

While *Wolbachia*-based interventions have shown promise in reducing vector-borne disease transmission, scaling these interventions for broad implementation in resource-limited settings remains challenging. A crucial aspect of this challenge is the need for efficient monitoring of *Wolbachia* introgression following the release of *w*Mel-infected *Ae. aegypti* mosquitoes. Effective surveillance of introgression and wildtype populations is essential for adjusting release strategies to ensure successful establishment. Our study demonstrates a strong correlation between *w*Mel prevalence estimates derived from ovitraps and those from BG-traps, supporting ovitraps as a reliable, cost-effective proxy for assessing *Wolbachia* introgression. Furthermore, our findings suggest that predictive models incorporating previous introgression data and mosquito abundance can further enhance the accuracy of ovitrap-based estimates. This evidence underscores the utility of ovitraps in ongoing monitoring and long-term assessment of W*olbachia* interventions, providing a scalable, low-resource tool that could facilitate widespread roll-out in low-income regions.

## Authors’ contributions

AIK, DATC, HS, LAM, EN, PMC conceived the project. EN and PMC conducted analyses and prepared the manuscript with supervision from GRdS, DATC, AIK. All authors reviewed, revised and approved the manuscript.

## Data and materials availability

All code and data used in these analyses are available at https://github.com/ko-laboratory/wmel-rj.

## Supplemental

**Fig S1. Significance analysis for oviposition trap larvae sample size.** The number of *Aedes aegypti* larvae samples needed at different levels of *w*Mel prevalence and significance levels are indicated by the colored lines, calculated using a two-sided power calculation for proportion test (one sample) from the ‘pwr’ package in R. The red horizontal line indicates the chosen sample size of 22 *Ae. aegypti* larvae. The blue dashed vertical line indicates 5% two-sided error.

**Table S1. BG-Sentinel trap sample collection and testing results.** Data is presented at the cluster level and combined across the three time points (March, April, May). Adult mosquitoes were trapped, morphologically-identified to the species level, and tested by WMP.

**Table S2. Oviposition trap sample collection and testing results.** Data is presented at the cluster level and combined across the three time points (March, April, May). Eggs were trapped and collected by the Belo Horizonte Municipal Health Department. Eggs were hatched, reared, and tested by the authors. Eggs were counted by the Municipality in March and April, and by an author in May.

**Table S3. BG-Sentinel trap unadjusted and adjusted prevalence estimates.** Estimates are calculated by averaging trap-level proportions within each cluster at each time point. The 95% binomial confidence interval is included after the prevalence point estimate.

**Table S4. Oviposition trap unadjusted and adjusted prevalence estimates.** Estimates are calculated by averaging trap-level proportions within each cluster at each time point. The 95% binomial confidence interval is included after the prevalence point estimate. Estimates marked as “NA” refer to time point without any *Aedes aegypti* larvae samples collected.

**Fig S2. Distribution of *w*Mel prevalence estimates in Ovitraps and BG-traps.** A) Histogram distribution of the BG-trap *w*Mel prevalence estimates by cluster and time point. B) Histogram distribution of the ovitrap *w*Mel prevalence estimates by cluster and time point.

**Table S5. Ovitrap introgression threshold classification sensitivity and specificity.** WMP uses 0.6 prevalence in an area as an introgression threshold to determine whether introgression will self-sustain or whether further releases are needed. The prevalence estimates from ovitraps were compared to those from BG-traps for all clusters and time points, and classification above or below the varying thresholds were used to construct 2×2 tables. Sensitivity and Specificity were calculated across clusters for each time point from the 2×2 tables.

**Fig S3. Within-Cluster *w*Mel Prevalence Variation.** The within-cluster variance of trap-level prevalence estimates was calculated and averaged by cluster and time point, along with the chi-square 95% confidence interval. The forest plots above show the distribution of the cluster-level variance by month. The plot was created using the *ggplot2* package in R. The variance across clusters for BG-traps was 0.11 in March, 0.13 in April, and 0.13 in May. For ovitraps it was 0.17 in March, 0.09 in April, and 0.11 in May.

**Table S6. Temporal correlation to inform variable selection for regression model.** Correlation estimates were calculated using a spearman-rank correlation test on each cluster’s *w*Mel count by time point. Given that only three time points were available for each cluster, there is not enough data for a significant p-value. Similarly, Pearson correlation tests were unable to run with the limited data.

**Table S7. Model predicted *w*Mel counts error.** Error was calculated by subtracting the true count from the model-predicted count and dividing by the true count. Negative error indicates an underestimation by the model, whereas positive error indicates an overestimation. NA resulted from a lack of ovitrap data at the time point or the previous time point. If 0 *w*Mel-positive were caught by BG-traps in that cluster at the time point, that estimate was excluded.

**Table S8. Model predicted introgression success classification sensitivity and specificity.** WMP uses 0.6 prevalence in an area as an introgression threshold to determine whether introgression will self-sustain or whether further releases are needed. The prevalence predicted by the models were compared to those from BG-traps across clusters and time points, and classification above or below the 0.6 threshold were used to construct 2×2 tables. Sensitivity and Specificity were calculated across clusters for each time point from the 2×2 tables.

**Table S9. Autocorrelation tests to determine spatial correlation of BG-traps.** Correlation estimates were calculated using a spearman-rank correlation test on each cluster’s *w*Mel count by time point. Given that only three time points were available for each cluster, there is not enough data for a significant p-value.

